# Pharmacokinetics and pharmacodynamics of inhaled antipseudomonal bacteriophage therapy in mice

**DOI:** 10.1101/2020.07.13.201798

**Authors:** Michael Y.T. Chow, Rachel Yoon Kyung Chang, Mengyu Li, Yuncheng Wang, Yu Lin, Sandra Morales, Andrew J McLachlan, Elizabeth Kutter, Jian Li, Hak-Kim Chan

## Abstract

Inhaled bacteriophage (phage) therapy is a potential alternative to conventional antibiotic therapy to combat multidrug-resistant (MDR) *Pseudomonas aeruginosa* infections. However, pharmacokinetics (PK) and pharmacodynamics (PD) of phages are fundamentally different to antibiotics and the lack of understanding potentially limits optimal dosing. The aim of this study was to investigate the *in vivo* PK and PD profiles of antipseudomonal phage PEV31 delivered by pulmonary route in mice. BALB/c mice were administered phage PEV31 at doses of 10^7^ and 10^9^ PFU by the intratracheal route. Mice (*n* = 4) were sacrificed at 0, 1, 2, 4, 8 and 24 h post-treatment and various tissues (lungs, kidney, spleen and liver), bronchoalveolar lavage and blood were collected for phage quantification. In a separate study, mice (*n* = 4) were treated with PEV31 (10^9^ PFU) or PBS at 2 h post-inoculation with MDR *P. aeruginosa*. Infective PEV31 and bacteria were enumerated from the lungs. In the phage only study, PEV31 titer gradually decreased in the lungs over 24 hours with a half-life of approximately 8 h for both doses. In the presence of bacteria, PEV31 titer increased by almost 2-log_10_ in the lungs at 16 h. Furthermore, bacterial growth was suppressed in the PEV31-treated group, while the PBS-treated group showed exponential growth. Some phage-resistant colonies were observed from the lung homogenates sampled at 24 h post-phage treatment. These colonies had a different antibiogram to the parent bacteria. This study provides evidence that pulmonary delivery of phage PEV31 in mice can reduce the MDR bacterial burden.

## Introduction

Since the discovery of penicillin, much effort has been targeted towards understanding the pharmacokinetics (PK) and pharmacodynamics (PD) of antibiotics to guide safe and effective treatment regimens. While bacteria can be intrinsically resistant to antibiotics, the inappropriate use of antibiotics has subjected bacteria to high selective pressure, leading to the advent of resistant strains at an alarming rate and now poses a serious global threat to human health. The severe threat of antimicrobial resistance remains imminent (1) and the World Health Organization has called for global action to tackle this crisis (2). Unfortunately, the antibiotic discovery pipeline is drying with a lack of novel antimicrobial agents against Gram-negative bacteria (3). In particular, the emergence of multidrug-resistant (MDR) *Pseudomonas aeruginosa* strains presents a major public health risk due to their prevalent intrinsic and acquired resistance to most antibiotics (4). MDR *P. aeruginosa* causes complications of respiratory infections associated with high morbidity and mortality rates in many diseases, including bronchiectasis, cystic fibrosis, chronic obstructive pulmonary disease and pneumonia (4).

Bacteriophages (phages) are naturally occurring bactericidal virus that infect targeted host bacteria. They are recently rediscovered and reintroduced as potential antimicrobial treatment and are considered an attractive solution to the increasing failure of antibiotics (5). Phage therapy predominantly relies on the lytic life cycle of phages. Virulent (lytic) phages recognize and attach to surface receptors of host bacterium, inject their genetic material and then utilize the metabolic machineries of the host for self-replication (5). Up to hundreds of progenies can be produced and then released into the surrounding during bacteriolysis. Phage therapy has distinct advantages over conventional antibiotic treatment in that phages are (i) a naturally occurring antibacterial, (ii) self-replicating, (iii) self-limiting upon resolution of infection, (iv) effective against both MDR or antibiotic sensitive bacteria, (v) highly specific with low inherent toxicity, (vi) able to co-evolve with bacteria, and (vii) able to penetrate biofilms (5). The potential use of phages as antibacterial agents has been demonstrated in *in vitro* (6, 7), preclinical (8–11) and in compassionate single case studies (12–14).

Despite these advantages and potential, development and application of phage therapy has been relatively slow. A possible reason is that the current understanding and paradigm associated with antibiotic treatment cannot be transferred directly to phages (15). The PK and PD of phages are fundamentally different from those of conventional antibiotics. While many antibiotics are small molecules, phages are nano-sized virus composed of proteins, nucleic acids (DNA or RNA) and sometimes lipids. In addition, phages have a unique dynamic with their bacterial host as self-replicating biopharmaceuticals (15). The PK/PD of phages are determined by their antibacterial activities featuring self-replication, phages and bacteria coevolution, as well as the human immune system in response to the two concurrent events (16).

Inhaled phage therapy holds remarkable potential to treat respiratory infections caused by bacteria, including MDR isolates (5). With oral inhalation route for delivery to the lung, high concentration of phages can be delivered to the site of infection in the respiratory tract, achieving high pulmonary bioavailability. Inhaled phage therapy has been used in Eastern European countries to treat bacterial respiratory infections that was otherwise untreatable with antibiotics. A 7-year old cystic fibrosis patient received inhaled phage therapy in 2011, which dramatically reduced the MDR *P. aeruginosa* and *Staphylococcus aureus* numbers in the lungs (12). Although inhaled phage therapy has been practiced in Eastern Europe for many decades, the phage viability in nebulized aerosol droplets has only recently been investigated. Our group and others have demonstrated that inhalable aerosolized *Pseudomonas* phages remain biologically active when a suitable nebulizer system is used (17–20). Although the feasibility of producing infective phage aerosols have been well-established, there is a lack of understanding of *in vivo* PK and PD profiles of phages and bacteria in the lungs. The aim of this study is to investigate the PK and PD profiles of *Pseudomonas* phage PEV31 administered by pulmonary delivery in a neutropenic murine model of respiratory infection.

## Materials and methods

### Bacteriophage

Anti-Pseudomonas phage PEV31 was isolated from the sewage plant in Olympia (WA, USA) by the Evergreen Phage Lab (Kutter Lab). PEV31 belongs to the *Podoviridae* family. Stocks of the phage were amplified using the Phage-on-Tap protocol (21) with minor modifications. Briefly, 200 mL of Nutrient Broth (NB, Amyl Media, Australia) supplemented with 1 mM of CaCl_2_ and MgCl_2_ were mixed with 0.1 volumes of overnight bacterial host (*P. aeruginosa* dog-ear strain PAV237). The mixture was incubated for 1 h with continuous shaking (220 rpm) at 37°C. A volume (200 μL) of PEV31 lysate at 10^9^ plaque forming units (PFU)/mL was added, followed by further incubation for 8 h. The mixture was centrifuged at 4000 × *g* for 20 min and the supernatant was filter-sterilized using 0.22 μm polyethersulfone membrane filter. The phage lysate was further purified and concentrated using ultrafiltration (100 kDa Amicon® Ultra-15 centrifugal filter, Sigma, Australia), and the media was replaced with phosphate-buffered saline (PBS) supplemented with 1 mM CaCl_2_. Bacterial endotoxins were removed by adding 0.4 volumes of 1-octanol, followed by vigorous shaking at room temperature for 1 h. The mixture was centrifuged at 4000 *g* for 10 min and then the aqueous phase was collected. Residual organic solvent was removed by centrifuging down the phages at 20,000 *g* for 1.5 h and then replacing the buffer with fresh PBS supplemented with 1 mM CaCl_2_.

### Endotoxin level quantification

The Endosafe^®^ Portable Test System (Charles River Laboratories, Boston, USA) was used as per the manufacturer’s instructions to quantify endotoxin level in the resulting phage lysates. The single use LAL assay cartridges contain four channels to which the LAL reagent and a chromogenic substrate have been pre-applied. A single cartridge enables duplicate measurements of the sample and positive control with a known endotoxin concentration. The sensitivity of the Readouts between 50% and 200% spike recovery are deemed acceptable. The sensitivity of the assay was 1 – 10 EU/mL. Endotoxin-free water, tips and tubes were used at all times.

### Bacterial strain and phage-susceptibility testing

*P. aeruginosa* FADDI-PA001 was used to induce bacterial infection in the mice used in this study. The strain is an MDR clinical isolate provided by Li lab, Monash University, Australia (8). Phage-susceptibility of this isolate was assessed using a spot test (6). Briefly, 5 mL of 0.4% Nutrient Broth top agar was mixed with overnight culture of FADDI-PA001 (approximately 2 × 10^8^ colony forming units, CFU) and then overlayed onto a 1.5% Nutrient Agar plate. Then, 10 μL of a phage stock solution was spotted on the top agar plate, air-dried and incubate at 37°C for 24 h. After incubation, the appearance of the lysis zone was assessed for phage-susceptibility.

### Animals

Female BALB/c mice of 6 – 8 weeks were obtained from Australian BioResources Ltd (Moss Vale, New South Wales, Australia). The mice were housed under a 12-hour dark-light cycle with *ad libitum* supply to standard chow diet and water. All animal experiments conducted were approved by the University of Sydney Animal Ethics Committee.

### Pharmacokinetics of PEV31 after intratracheal administration

Healthy (non-infected) mice were anaesthetized by intraperitoneal injection of ketamine / xylazine mixture (80 mg/kg and 10 mg/kg, respectively) in 150 μl PBS. Upon deep anesthesia as confirmed by the absence of pedal reflex, the anaesthetized mouse was suspended on a nylon floss by its incisor teeth and placed on an inclined intubation board. The trachea was then gently intubated with a soft plastic guiding cannula. PEV31 at two different doses (10^7^ and 10^9^ PFU) suspended in 25 μL PBS was administered into the trachea through the guiding cannula with a micropipette and a 200 μL gel-loading pipette tip. At 0 (immediately after administration), 1, 2, 4, 8 and 24 h post-phage administration, mice (*n* ≥ 4) were terminally anaesthetized by intraperitoneal injection of an overdose of ketamine / xylazine mixture (300 mg/kg and 30 kg/mg, respectively). Broncho-alveolar lavage (BAL), lung, kidney, spleen, liver and blood samples were collected (**Figure 1A**). BAL was performed by instilling 1.5 mL PBS (as three aliquots of 0.5 mL) through the trachea to the lung and collecting the lavage suspension. Approximately 1.2 to 1.3mL of lavage suspension was recovered. Harvested tissues were homogenized by TissueRuptor II with plastic probes (QIAGEN, Hilden, Germany) in cold PBS under ice-water bath for 30 seconds. Tissue homogenates were kept at 2 – 8 °C until phage quantification by plaque assay (described below). Plaque assay was performed within 3 hours of sample collection.

**Figure 1.**
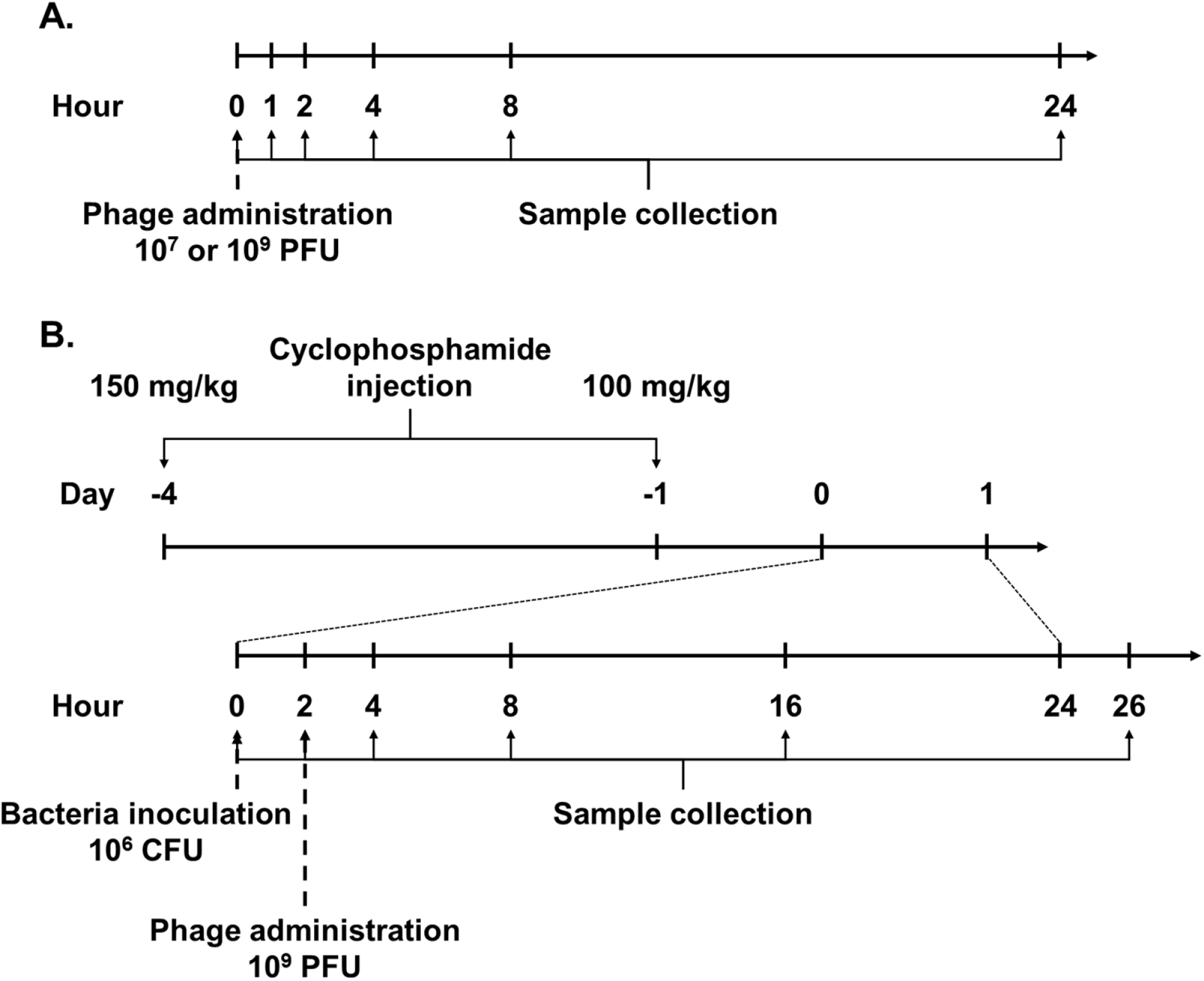
Timeline of experimental procedures to investigate the pharmacokinetics (A) and pharmacodynamics (B) of intratracheally administered PEV31 in mice.

### Pseudomonas pulmonary infection

A neutropenic murine model (8) was used to establish pulmonary *P. aeruginosa* infection. Two doses of cyclophosphamide were intraperitoneally administered 4 days (150 mg/kg) and 1 day (100 mg/kg) prior to infection. On the day of infection, the FADDI-001 bacterial suspension at its early logarithmic growth phase was inoculated intratracheally at a concentration of 10^6^ CFU in 25 μL, as described above. At 2 h post-infection, PEV31 suspension (10^9^ PFU in 25 μL PBS) or sterile PBS of equal volume (as untreated control) was intratracheally administered to the infected mice. Following terminal anesthesia as described above, BAL and other tissues were collected at 0 (immediately after bacteria inoculation), 2 (immediately after phage administration), 4, 8, 16 and 26 h post-infection (*n* = 4) (**Figure 1B)**. Collected tissues were homogenized in cold PBS. Bacterial load and phage concentrations in the tissue homogenates and BAL were stored on ice at all times and analyzed within two hours using plaque assay and colony counting, respectively, as described below. Bacteria enumeration was performed as soon as practically possible and was not later than 2 hours after sample collections. Plaque assay was performed within 3 hours of sample collection.

### Plaque assay

BAL and tissue homogenates were serially diluted in PBS for phage quantification. For samples from infected animals, bacteria were first removed from the homogenates and BAL samples by filtering through 0.22 μm polyethersulfone membrane filter before dilution. A volume of reference bacterial host (PAV237) containing 2 × 10^8^ CFU at stationary phase was mixed with 5 mL of Nutrient Broth top agar. The mixture was overlaid on top of a Nutrient Agar plate and dried for 15 min. Then, 20 μL of serially diluted phage suspension were dropped on top of the top agar plate, left to air dry, and then incubated for 24 h at 37°C. The diluted samples were analyzed in triplicate.

### Bacteria enumeration

Phage-inactivation was performed prior to bacteria enumeration to prevent bactericidal activities of phages and impede reduction of CFU counts *ex vivo*. Furthermore, all samples were kept on ice at all times to minimize the risk of phage-bacteria interactions *in vitro*. The samples were treated with tannic acid (20 mg/L) and ferrous sulfate (2.5 mM) to inactivate the phage PEV31 and then treated with 2% Tween 80 in PBS to stop the interaction. The phage-inactivated lung homogenate was filtered through a sterile filter bag (bag stomacher filter with a pore size of 280 μm, Labtek Pty Ltd., Australia). Filtrate samples and BAL were serially diluted in PBS and then spiral plated on Nutrient Agar plates using an automatic spiral plater (WASP, Don Whitley Scientific, United Kingdom). The plates were air-dried and then incubated for 24 h at 37°C. Colonies were counted using a ProtoCOL automated colony counter (Synbiosis, United Kingdom).

### Minimum inhibitory concentration

Bacterial colonies from the spiral plates (*t* = 0 and 26 h) were taken and inoculated in Nutrient Broth. The antibiogram of these colonies was assessed by determining the minimum inhibitory concentrations (22) of selected antibiotics, including ciprofloxacin, tobramycin, colistin and aztreonam. A volume (190 μL) of early-log phase bacterial culture (1 × 10^6^ CFU/mL) was mixed with 10 μL of antibiotics (0.25, 0.5, 1, 2, 4, 8, 16, 32, 64 μg/mL). The treated bacterial culture was incubated for 24 h at 37°C with continuous shaking at 220 rpm. Optical density at 600 nm (OD_600_) was measured using a microplate reader (Victor multilabel Plate Reader, Perkin Elmer, United States).

### Cytokine quantification

The BAL collected was centrifuged at 400 × *g* for 10 min, and the supernatant was collected as the broncho-alveolar lavage fluid (BALF). The levels of interleukin (IL)-6, TNF-alpha and IL-1β (in infected mice) in BALF were quantified by enzyme-linked immunosorbent assay (ELISA) according to the manufacturer’s protocol (DY406, DY410 and DY401 from R&D systems; Minnesota, USA). The UV absorbance at 450 nm and 570 nm were measured for the primary signal and for plate correction, respectively (Victor multilabel Plate Reader). The standard curves were constructed by 4-parameter logistic non-linear regression.

### Data analysis

For the pharmacokinetics study, regression analysis on the phage titer (PFU) in lungs (lung tissues and BALF combined) over time was performed using simple exponential decay model. The exponential decay takes the form of *P*_*t*_ = *P*_0_ × *e*^−kt^ where *P*_*t*_ and *k* are the relative phage titer at time *t* and the rate constant (in h^-1^) respectively (*P*_0_ denoted the phage titer at time = 0). It follows that the half-life of the decay was given by *ln*2/*k*. The regression was performed using Prism software version 8.3 for Windows (GraphPad Software Inc., California, USA). The regressions were done with the sums of the squares weighted by the reciprocal of the dependent variables squared (*i.e.* 1/*y*^2^).

## Results

### Phage preparation and in vitro phage susceptibility

The titer of purified PEV31 enumerated against the reference strain (PAV237) used for phage amplification was 4 × 10^10^ PFU/mL. Phage PEV31 formed a clear zone of lysis on top of overlay plate containing MDR *P. aeruginosa* isolate FADDI-001. PEV31 was highly efficacious *in vitro* against FADDI-001 with an efficiency of plating of 1. Endotoxin level in the purified phage lysate was 3.8 EU/mL (i.e. 0.095 EU in 25 μL). The spike recovery was 130% and the coefficient of variation of the assay was 4%, which were all considered acceptable as per the manufacturer’s recommendations.

### Pharmacokinetics of intratracheally administered PEV31

Infectious PEV31 gradually decreased in the lungs (lung homogenate and BALF combined) over time regardless of the administered dose (**Figure 2**). At 24 h after IT administration, the phage titer dropped to 12.3% and 15.2% of the administered dose for the low (10^7^ PFU) and high (10^9^ PFU) doses, respectively. The elimination of active phage could be adequately described using simple exponential decay model, with the adjusted weighted *R*^2^ being 0.864 and 0.956 for the low and high dose, respectively. The rate constant *k* was estimated to be 0.0875 h^-1^ (95% CI 0.0527 to 0.0924) for the low dose and 0.0797 h^-1^ (95% CI 0.0700 to 0.0871) for the high dose, which are equivalent to a half-life of 7.9 (95% CI 7.5 to 13.2) and 8.7 (95% CI 8.0 to 9.9) hours, respectively. Infective PEV31 titer in other organs including kidney, liver, blood and spleen was extremely low and only accounted for less than 0.01% of the administered doses (**Figure 3**). The phage titer gradually increased over 24 hours in the liver of mice that received 10^9^ PFU of PEV31. PEV31 suspension was well tolerated at a low dose without changes in inflammatory cytokine level (**Figure 4**). On the contrary, a transient upregulation of TNF-α and IL-6 activity was observed at 4 and 8 h post-administration, respectively when the mice were given a high dose of PEV31. Both cytokines returned to baseline at 24 h after the single IT dose.

**Figure 2.**
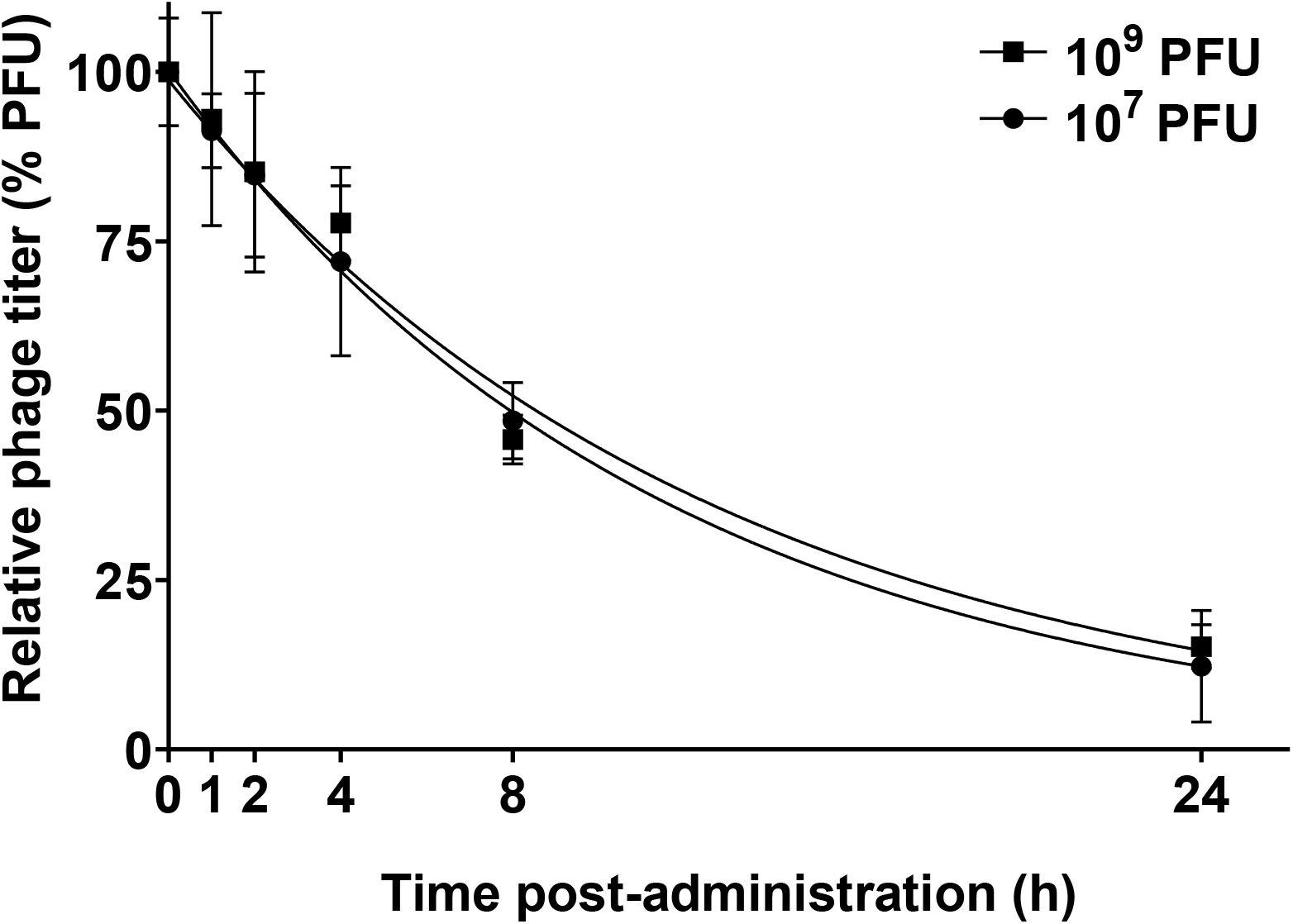
Relative phage titer in the lungs (lung tissues and BALF combined) of healthy mice after intratracheal administration of phage PEV31 at doses of 10^7^ and 10^9^ PFU. Phage titer is expressed as number of PFU relative to the administered dose. Error bar denotes standard deviation (*n* ≥ 4 except for *t* = 2 h of the 10^7^ PFU group, and *t* = 1 h and 4 h of the 10^9^ PFU group where *n* = 3).

**Figure 3.**
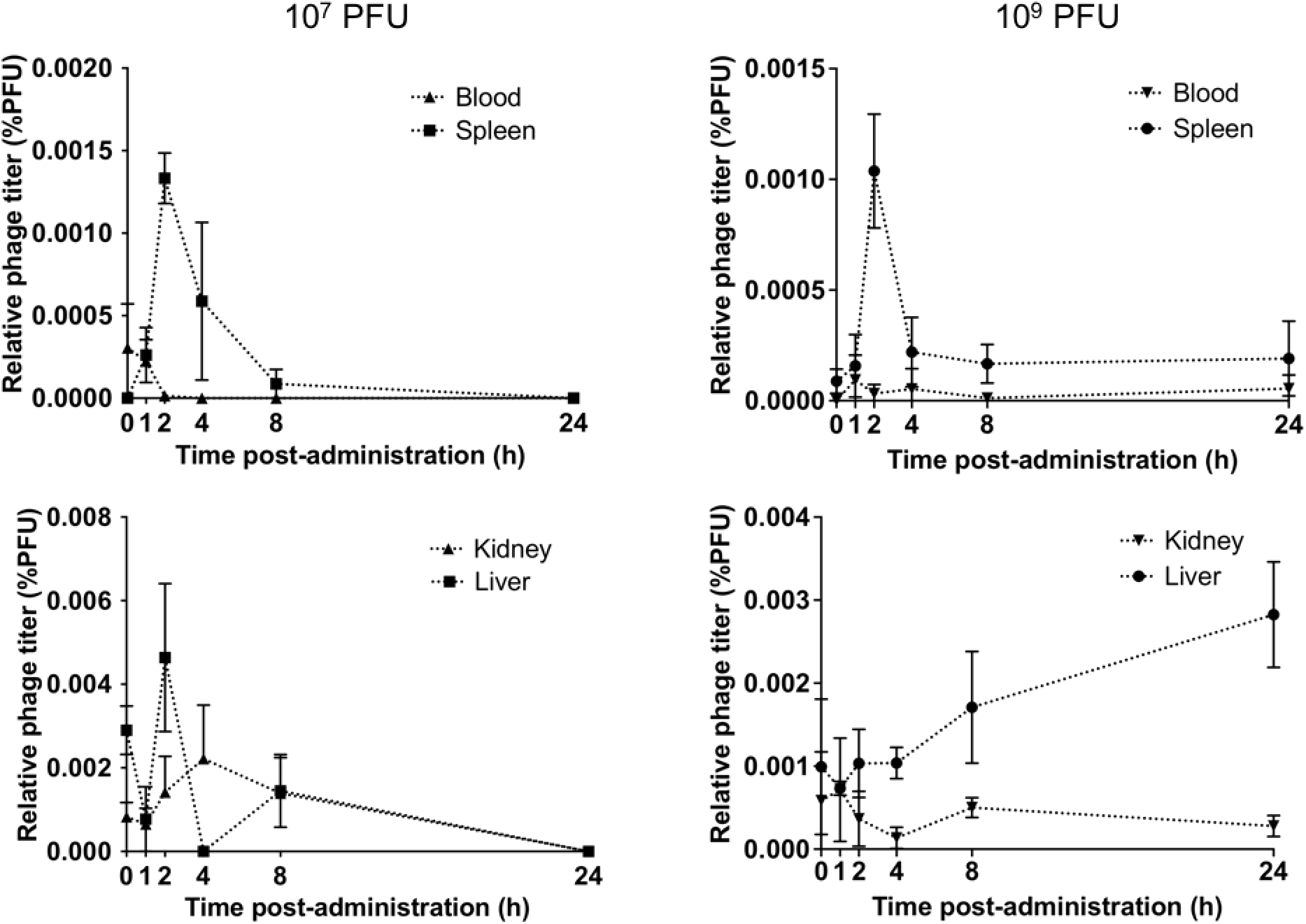
Relative phage titer in kidney, liver, blood and spleen at two phage doses. Phage titer is expressed as number of PFU relative to the administered dose. Error bar denotes standard error (n ≥ 4).

**Figure 4.**
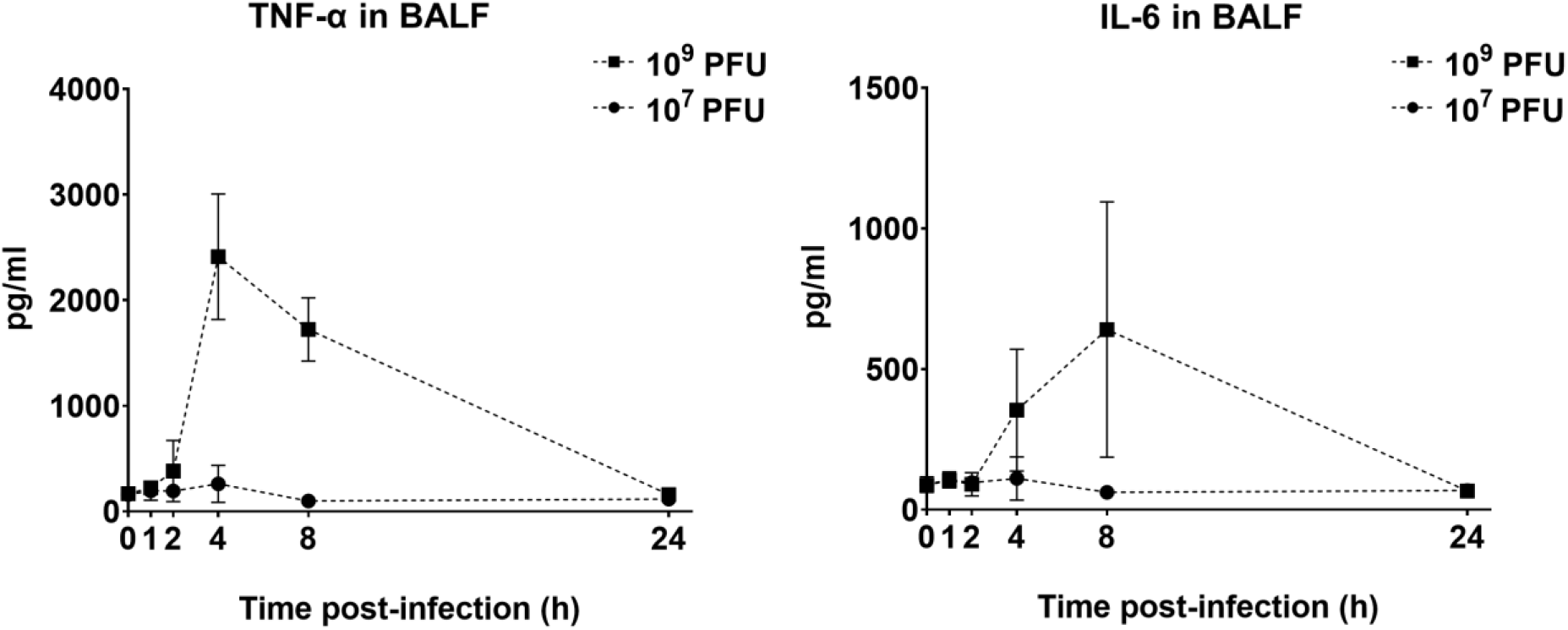
Levels of TNF-α and IL-6 in BALF in healthy mice after phage administration at doses of 10^7^ and 10^9^ PFU. Error bar denotes standard deviation (*n* ≥ 4 except for *t* = 2 h of the 10^7^ PFU group where *n* = 3).

### Pharmacodynamics of intratracheally administered PEV31

In the infected only group, the bacteria continued to replicate without any significant stationary period (**Figure 5**). Initially, the bacteria grew exponentially for up to about 8 h post-infection (hpi), after which the growing rate decreased. In the infected mice treated with 10^9^ PFU of PEV31 at 2 hpi, the bacterial load in lung remained mostly unchanged except for the initial drop at 4 hpi (2 h post-phage IT administration), conferring to more than 4-log reduction in bacterial load at 26 hpi. Some of the survived bacterial colonies at 26 hpi in the phage-treated group showed a different antibiogram profile in comparison with the parent bacteria used to inoculate each mouse (**Table 1**). The MIC value of ciprofloxacin decreased from 8 to 2 μg/mL, while tobramycin and colistin increased from 8 to 64 μg/mL and 4 to 32 μg/mL, respectively. There were no apparent sensitivity changes to other antibiotics tested and all the tested bacterial colonies from PBS-treated group had the same MIC as the parent stock.

**Table 1.** Antibiotic and phage susceptibility of bacterial colonies isolated from the lung homogenate before and after treatment with phage PEV31 or PBS.

| Time post-infection (h) | Treatment | Number of colonies observed (tested) | MIC (µg/mL) |  |  |  |  | PEV31 susceptibility |
| --- | --- | --- | --- | --- | --- | --- | --- | --- |
|  |  |  | Amikacin | Ciprofloxacin | Tobramycin | Aztreonam | Colistin |  |
| 0 | n/a | 4 (4) | 32 | 8 | 8 | >64 | 4 | S |
| 26 | PBS | 4 (4) | 32 | 8 | 8 | >64 | 4 | S |
| 26 | Phage | 3 (8) | 32 | 2 | 64 | >64 | 32 | R |
| 26 | Phage | 5 (8) | 32 | 8 | 8 | >64 | 4 | S |

NOTE: MIC, minimum inhibitory concentration; S, susceptible; R, resistant.

**Figure 5.**
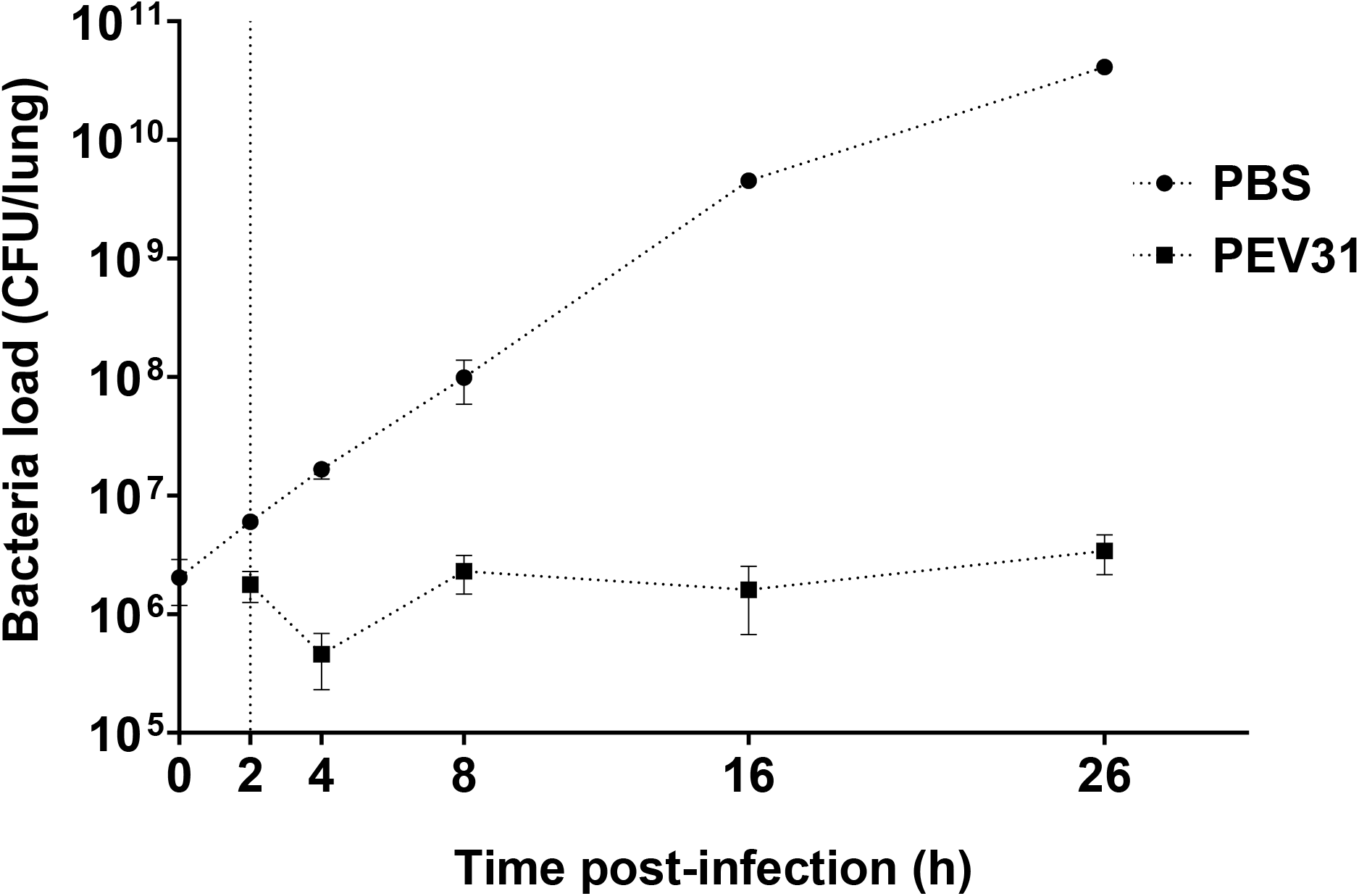
Bacterial load in the lungs of mice treated with PBS and phage over 26 hours. Dotted vertical line represents time of phage administration (*t* = 2 h). Error bar denotes standard deviation (*n* = 2 – 4).

The phage-mediated bacterial killing was evident by increase of infectious phage particles over time (around 2-log_10_) at 24 h post-phage administration (**Figure 6**). Inflammatory cytokines activity (TNF-α, IL-1β and IL-6) in BALF were also measured as an evaluation of lung inflammation. In bacteria-infected only group, a substantial upregulation of all three cytokines was observed. TNF-α peaked at 4 hpi while the other two cytokines displayed peak activity at later time points. The upregulation of IL-1β activity considerably diminished at 26 hpi and to a lesser extent for TNF-α, but not for IL-6. Interestingly, the upregulation in cytokines was only partially suppressed by the phage treatment for IL1-β (23), but not for TNF-α and IL-6. In the phage-treated group, the peak of TNF-α appeared delayed to 8 hpi.

**Figure 6.**
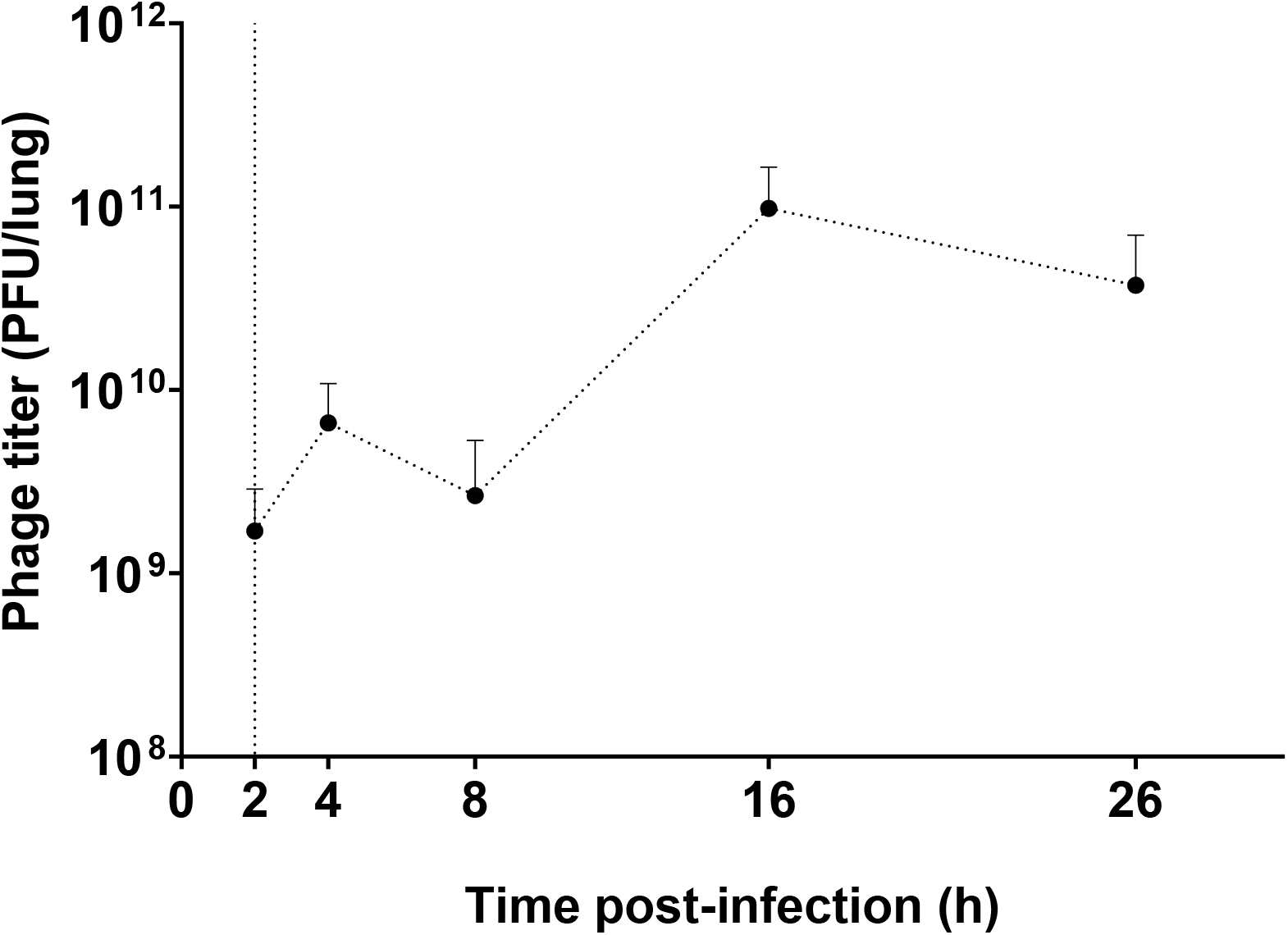
Phage titer in the lungs of mice infected with *P. aeruginosa*. Dotted vertical line represents the time (*t* = 2 h) of phage administration (10^9^ PFU). Error bar denotes standard deviation (*n* = 3 – 4).

## Discussion

This is the first study investigating PK and PD of intratracheally administered *Pseudomonas* phages *in vivo*. In previous studies with mice, the intranasal route has been widely used for initiation of lung infection and then phage treatment (24–26) likely due to ease of administration. These studies provide strong support for inhaled phage therapy with reduction in bacterial load and inflammation in the mouse lung infection model. Compared with intranasal route, intratracheal administration enables direct application of bacteria and phages to the mouse lungs with minimal loss in other parts of the respiratory route, including nose, throat and upper airways (8, 27). Hence, the exact phage doses of interest were given in the PK study, and in the PD study. Despite these advantages, studies on intratracheal administration of phages for lung infections have been scarce (8, 28). In this study, intratracheal instillation was used to administer and assess the PK of phage PEV31 at two different doses.

The infectious phage PEV31 as quantified using plaque assay exhibited an exponential decay in the lung at both low and high doses with similar half-life (rate constant, **Figure 2**). After oral inhalation of phages to the lung, the phage titer dropped by 1-log_10_ over 24 hours at both doses. Liu *et al.* studied the PK profile of *Siphoviridae* lytic mycobacteriophage D29 after doing 5 × 10^8^ PFU via intra-tracheal route (28). The titer of D29 dropped to 1.2-log_10_ by 24 hours post-administration. Using the same regression methodology on the titers reported by Liu *et al*., we determined the half-life of D29 to be 5.8 hours, which is lower than our values of 7.9 – 8.7 hours for PEV31. Phage D29 belongs to the *Siphoviridae* family and has a longer phage tail as compared with PEV31 (*Podoviridae*). Whether there is a correlation between the family and/or the geometry of the phage particle and the rate of elimination warrants further investigations. Compared with phage delivered via intravenous injection (10), intra-tracheal route resulted in reduced systemic exposure (**Figure 3**).

Our current work has shown that the total titer of administered PEV31 phages in various organs do not add up to 100% of the delivered dose. No phage titer reduction was observed during the sample processing, including homogenization, filtration (0.22 μm PES membrane and BagPage filter) and sample dilution. This implies that phage inactivation or degradation in the lungs and/or other organs are likely. Hence, both biodistribution of phages as well as degradation/inactivation may contribute to the titer reduction observed in the lungs over time. Plaque assay is the method of choice for quantifying infectious phages (29, 30). In a plaque assay, a zone of clearance (plaques) are formed on top of a bacterial lawn as a result of cycles of infection of the bacterial cells with phage progeny radiating from the original source of infection (31). It follows that only infectious phages can be enumerated. To evaluate the total number of viral particles – infectious, non-infectious and defective, genome quantification using qPCR can potentially be utilized (32, 33). The combination of qPCR and plaque assay could potentially help understand the biodistribution of infective phages as well as those that have been broken down or inactivated in different organs. This information may be particularly useful for estimating the total phage burden over time and correlating any long-term side effects associated with accumulation of nano-sized virus particles in the various cavities during a prophylactic treatment. Nano-sized particles (<10 nm) can easily enter human tissues and disrupt the biochemical environment of normal cells (34–38). Nanoparticles mostly accumulate in the liver tissues and adverse unpredictable health outcomes have recently surfaced (39, 40). Our current study has shown that infective phage titer gradually increases in the liver over time, and there may be inactive or degraded phage particles further accumulated in the organ. The most significant phage phagocytosis function is thought to be played by the liver. Phages mostly accumulate in the liver (99%) after intravenous administration and the rate of phagocytosis by Kupffer cells are four times faster than splenic macrophages (41). The current study has shown that by delivering phages directly to the lungs, systemic exposure and liver-induced phagocytosis is substantially minimized.

In the current study, phage PEV31 at a high dose [0.095 EU; 4.75 EU/kg] resulted in an upregulation of the inflammatory cytokine TNF-α at 2 h post-administration, which then substantially increased at 4 h (**Figure 4**). The upregulation of IL-6 followed and then peaked at a later timepoint of 8 h. Upon phage administration, TNF-α and other inflammatory cytokines (e.g. IL-1) secreted from resident macrophages stimulated the release of other chemoattractant factors such as MIP-2, MCP-1 and IL-6, promoting the adherence of circulating inflammatory cells to the endothelium (28). The upregulation of both cytokines subsided at 24 h. Overexpression of these cytokines were absent when the mice were administered a lower dose. It has been reported that phages could trigger both inflammatory and anti-inflammatory responses (42), and endotoxin alone cannot explain all the observed upregulation of cytokines in the current study. Liu *et al.* reported no significant differences in leukocyte, neutrophils, lymphocytes and TNF-α levels in the BALF at 24 h post-administration of D29 in healthy mice (28). However, the levels of these cytokines between 0 and 24 h post-administration is unknown and unfortunately, the endotoxin level in the phage preparations was unreported. In another study, intra-nasal administration of phages with endotoxin level of 0.0063 EU/mice (approx. 0.3 EU/kg) did not exhibit appreciable levels of TNF-α at 48 h post-treatment (9). Extremely low TNF-α and IL-6 levels were similarly observed in the lungs of mice that received *Pseudomonas* phages via intra-nasal route at 24 h post-administration, although the endotoxin level of the phage lysate was unreported (24). These findings aligned with our observation, where the upregulation of inflammatory cytokines subsided at 24 h post-administration. Hence, phage preparations with endotoxin levels even lower than that for parenteral and free of bacterial impurities should be considered for respiratory delivery (42) to ensure minimum toxicity (43), particularly in the case of prophylactic use. The current consensus is that phage therapy is safe (and has been so for decades) provided the phage preparation is sufficiently purified with low endotoxin level and other bacterial impurities (5, 44, 45). However, phage lysates originated from Gram-negative pathogens can be contaminated with endotoxins (lipopolysaccharides) and other proteins that are toxic to humans. Endotoxins are highly immunogenic and may cause septic shock by triggering cytokine signaling (46–48). The highest permitted endotoxin concentration for injection is 5 EU/kg/h. Even purified phage preparations with extremely low endotoxin level (<0.1 EU) may induce some pro-inflammatory responses, which are caused by other bacterial proteins and nucleic acids present in the phage lysate (42). Hence, despite low endotoxin level in our PEV31 preparation, bacterial proteins and other contaminants may have caused pro-inflammatory response in the murine lungs. Lung infection with *P. aeruginosa* caused significant increase in inflammatory cytokines (TNF-α, IL-1β and IL-6), with levels of IL-1β partially suppressed by phage treatment (**Figure 7**). Immune responses of phages are phage specific and some phages can even be anti-inflammatory such that bacteria clearance is reduced to promote phage propagation, as well as minimize the clearance of phages from the site of infection (42). Anti-inflammatory cytokines (e.g. IL-10) can be investigated in future studies to assess the potential role of phages as an anti-inflammatory agent.

**Figure 7.**
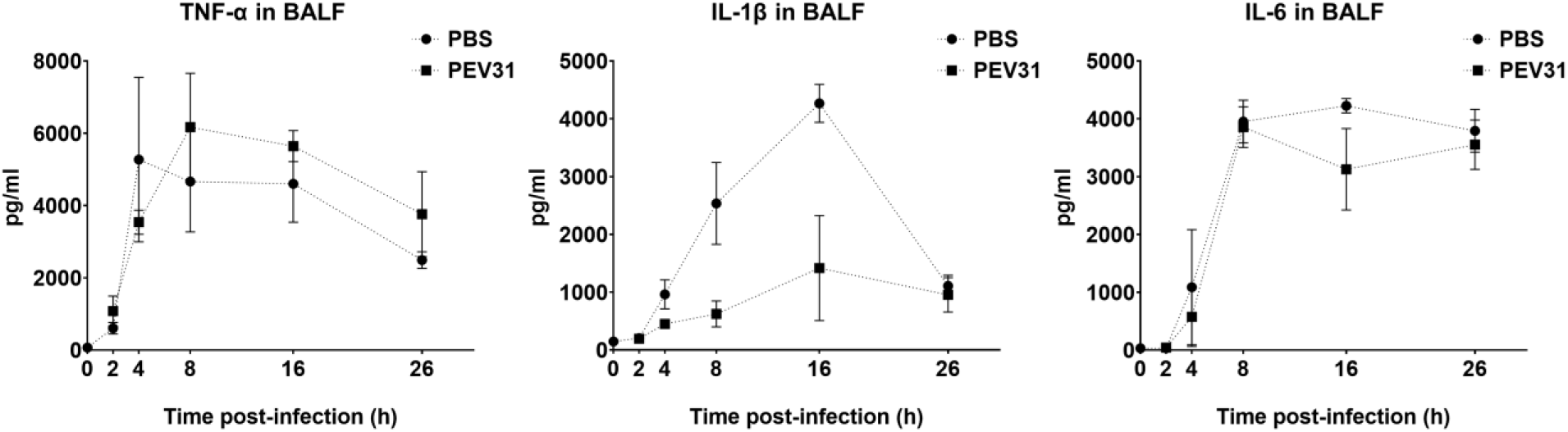
Levels of IL-6, TNF-α and IL-1β in BALF of *P. aeruginosa*-infected mice treated with PBS or phage PEV31 (10^9^ PFU) over time. Error bar denotes standard deviation (*n* = 2 – 4).

MDR *P. aeruginosa* isolate FADDI-001 used in this study is a clinical strain isolated from an ICU patient and is resistant to multiple antibiotics including rifampicin, doxycycline, ciprofloxacin, amikacin, aztreonam and tobramycin. When PEV31 was administered two hours post infection, the growth of FADDI-001 was suppressed, suggesting the rate of bacteria replication was similar to that of bacteria removal through phage-killing or clearance by host immune response, with the former evident by the increase in phage titer (**Figure 6**). FADDI-001 was highly susceptible to PEV31 *in vitro* but developed phage-resistance over time (upon overnight incubation) as reported for many naturally occurring phages under *in vitro* conditions. Bacteria can resist phage infection through different mechanisms, including (i) spontaneous mutations to prevent phage adsorption or phage DNA entry, (ii) restriction modification systems to cut phage nucleic acids, and (iii) CRISPR-Cas system mediated adaptive immunity (49–53). When bacteria are pressured with a high number of infective phages, resistance can develop rapidly (54) by changing the bacterial surface components that act as phage-binding receptors. These receptors can be blocked by producing extracellular matrix or competitive inhibitors, or even be removed to prevent phage adsorption (49). Contrary to *in vitro* results (data not shown), some but not all bacteria at 24 hpi remained susceptible to the phage despite high initial MOI. This may be due to fundamental differences between *in vitro* and *in vivo* systems, such as the involvement of mammalian immune responses and heterogeneous mixing. For the latter, it is possible that bacteria and phages were not evenly mixed within the mouse lungs during administration (*i.e.* spatial constrain). Hence, not all the bacteria may have been exposed to the same stress and selective pressure despite high phage titer used in this study. Those colonies that became resistant to phage PEV31 showed a different antibiogram to phage-susceptible bacteria. In the fight to become phage-resistance, FADDI-001 developed increased sensitivity to ciprofloxacin (quinolone), but also developed increased resistance to tobramycin (aminoglycoside) and colistin (polymyxin) (Table 1). The changes to antibiogram suggests possible modifications in the bacterial cell envelope as a result of acquiring phage resistance. The susceptibility of amikacin (another aminoglycoside) remained the same, suggesting the phage-mediated mechanisms for the PEV31-FADDI-001 system to antibiotic susceptibility is antibiotic-specific. Reversal of antibiotic resistance of *P. aeruginosa* under selective pressure of phage has been reported (55). Chan *et al.* isolated a lytic *Pseudomonas* phage OMKO1 that binds to outer membrane porin M of the multidrug efflux systems. MDR *P. aeruginosa* developed resistance to OMKO1 within 24 h of incubation *in vitro*, while these phage-resistant bacteria regained sensitivity to ciprofloxacin, erythromycin, ceftazidime and tetracycline. Understanding the phage-mediated mechanisms to antibiotic susceptibility is outside the scope of this work. However, the data suggested the need to assess the impact of phage treatment on antibiotic susceptibilities for each phage-bacteria system, particularly if combined phage-antibiotic treatment is being considered in a clinical setting.

The current study used an acute lung infection mouse model and does not necessarily inform the PK and PD data in chronic infections, such as cystic fibrosis. One of the major challenges in conducting simultaneous PK/PD study of phage therapy in bacterial infection lies in the continued interactions between bacterial host and phage even after sample collection. Procedures have been taken to minimize lysis of bacteria and phage propagation once the lung tissues have been harvested through physical separation by filtration and chemical inactivation by viricides. However, complete elimination of phage from bacteria in the tissue homogenates and removing phages that have already infected the bacteria could be difficult. Any remaining phages that have not been removed or inactivated could reduce the bacteria count and thus overestimate phage killing efficacy.

## Conclusion

This is the first study investigating the PK and PD profiles of antipseudomonal phage in the lungs of healthy and *P. aeruginosa*-infected mice, respectively. The safety and biodistribution of phage PEV31 over time were assessed in the lungs of healthy mice. Importantly, inhaled phages not only reduced the lung bacterial load, but also suppressed pro-inflammatory cytokines in the lungs. Bacterial antibiogram was altered upon phage treatment, where bacteria became susceptible to some, and more resistant to other antibiotics. Nonetheless, more work is required to examine the influence of phage exposure on antibiotic susceptibility of bacteria. Further *in vivo* toxicity and PK/PD studies evaluating various dose regimes in both acute and chronic models are urgently needed to better understand the phage and bacteria kinetics in the lungs.

## Acknowledgement

This study was financially supported by National Health and Medical Research Council (Project Grant APP1140617).

## References

1. CDC. 2019. Antibiotic Resistance Threats in the United States, 2019, *on* U.S. Department of Health and Human Services, CDC. Accessed 5th June.

2. WHO. 2015. Global action plan on antibicrobial resistance. https://www.who.int/antimicrobial-resistance/publications/global-action-plan/en/. Accessed

3. Theuretzbacher U, Outterson K, Engel A, Karlen A. 2020. The global preclinical antibacterial pipeline. Nat Rev Microbiol 18:275–285.

4. Pang Z, Raudonis R, Glick BR, Lin TJ, Cheng Z. 2019. Antibiotic resistance in Pseudomonas aeruginosa: mechanisms and alternative therapeutic strategies. Biotechnol Adv 37:177–192.

5. Chang RYK, Wallin M, Lin Y, Leung SSY, Wang H, Morales S, Chan HK. 2018. Phage therapy for respiratory infections. Adv Drug Deliv Rev 133:76–86.

6. Chang RY, Wong J, Mathai A, Morales S, Kutter E, Britton W, Li J, Chan HK. 2017. Production of highly stable spray dried phage formulations for treatment of Pseudomonas aeruginosa lung infection. Eur J Pharm Biopharm 121:1–13.

7. Pallavali RR, Degati VL, Lomada D, Reddy MC, Durbaka VRP. 2017. Isolation and in vitro evaluation of bacteriophages against MDR-bacterial isolates from septic wound infections. PLoS One 12:e0179245.

8. Chang RYK, Chen K, Wang J, Wallin M, Britton W, Morales S, Kutter E, Li J, Chan HK. 2018. Proof-of-principle study in a murine lung infection model of Antipseudomonal activity of phage PEV20 in a dry-powder formulation. Antimicrob Agents Chemother 62.

9. Carmody LA, Gill JJ, Summer EJ, Sajjan US, Gonzalez CF, Young RF, LiPuma JJ. 2010. Efficacy of bacteriophage therapy in a model of Burkholderia cenocepacia pulmonary infection. J Infect Dis 201:264–71.

10. Lin YW, Chang RY, Rao GG, Jermain B, Han ML, Zhao JX, Chen K, Wang JP, Barr JJ, Schooley RT, Kutter E, Chan HK, Li J. 2020. Pharmacokinetics/pharmacodynamics of antipseudomonal bacteriophage therapy in rats: a proof-of-concept study. Clin Microbiol Infect doi:10.1016/j.cmi.2020.04.039.

11. Wang JL, Kuo CF, Yeh CM, Chen JR, Cheng MF, Hung CH. 2018. Efficacy of phikm18p phage therapy in a murine model of extensively drug-resistant Acinetobacter baumannii infection. Infect Drug Resist 11:2301–2310.

12. Kvachadze L, Balarjishvili N, Meskhi T, Tevdoradze E, Skhirtladze N, Pataridze T, Adamia R, Topuria T, Kutter E, Rohde C, Kutateladze M. 2011. Evaluation of lytic activity of staphylococcal bacteriophage Sb-1 against freshly isolated clinical pathogens. Microb Biotechnol 4:643–50.

13. Zhukov-Verezhnikov NN, Peremitina LD, Berillo EA, Komissarov VP, Bardymov VM. 1978. [Therapeutic effect of bacteriophage preparations in the complex treatment of suppurative surgical diseases]. Sov Med:64–6.

14. Ujmajuridze A, Chanishvili N, Goderdzishvili M, Leitner L, Mehnert U, Chkhotua A, Kessler TM, Sybesma W. 2018. Adapted Bacteriophages for Treating Urinary Tract Infections. Front Microbiol 9:1832.

15. Principi N, Silvestri E, Esposito S. 2019. Advantages and Limitations of Bacteriophages for the Treatment of Bacterial Infections. Front Pharmacol 10:513.

16. Roach DR, Leung CY, Henry M, Morello E, Singh D, Di Santo JP, Weitz JS, Debarbieux L. 2017. Synergy between the Host Immune System and Bacteriophage Is Essential for Successful Phage Therapy against an Acute Respiratory Pathogen. Cell Host Microbe 22:38–47 e4.

17. Leung SSY, Carrigy NB, Vehring R, Finlay WH, Morales S, Carter EA, Britton WJ, Kutter E, Chan HK. 2019. Jet nebulization of bacteriophages with different tail morphologies - Structural effects. Int J Pharm 554:322–326.

18. Carrigy NB, Chang RY, Leung SSY, Harrison M, Petrova Z, Pope WH, Hatfull GF, Britton WJ, Chan HK, Sauvageau D, Finlay WH, Vehring R. 2017. Anti-Tuberculosis Bacteriophage D29 Delivery with a Vibrating Mesh Nebulizer, Jet Nebulizer, and Soft Mist Inhaler. Pharm Res 34:2084–2096.

19. Astudillo A, Leung SSY, Kutter E, Morales S, Chan HK. 2018. Nebulization effects on structural stability of bacteriophage PEV 44. Eur J Pharm Biopharm 125:124–130.

20. Marqus S, Lee L, Istivan T, Kyung Chang RY, Dekiwadia C, Chan HK, Yeo LY. 2020. High frequency acoustic nebulization for pulmonary delivery of antibiotic alternatives against Staphylococcus aureus. Eur J Pharm Biopharm 151:181–188.

21. Bonilla N, Rojas MI, Netto Flores Cruz G, Hung SH, Rohwer F, Barr JJ. 2016. Phage on tap-a quick and efficient protocol for the preparation of bacteriophage laboratory stocks. PeerJ 4:e2261.

22. Chang RYK, Das T, Manos J, Kutter E, Morales S, Chan HK. 2019. Bacteriophage PEV20 and Ciprofloxacin Combination Treatment Enhances Removal of Pseudomonas aeruginosa Biofilm Isolated from Cystic Fibrosis and Wound Patients. AAPS J 21:49.

23. Dinarello CA. 2018. Overview of the IL-1 family in innate inflammation and acquired immunity. Immunol Rev 281:8–27.

24. Debarbieux L, Leduc D, Maura D, Morello E, Criscuolo A, Grossi O, Balloy V, Touqui L. 2010. Bacteriophages can treat and prevent Pseudomonas aeruginosa lung infections. J Infect Dis 201:1096–104.

25. Morello E, Saussereau E, Maura D, Huerre M, Touqui L, Debarbieux L. 2011. Pulmonary bacteriophage therapy on Pseudomonas aeruginosa cystic fibrosis strains: first steps towards treatment and prevention. PLoS One 6:e16963.

26. Alemayehu D, Casey PG, McAuliffe O, Guinane CM, Martin JG, Shanahan F, Coffey A, Ross RP, Hill C. 2012. Bacteriophages phiMR299-2 and phiNH-4 can eliminate Pseudomonas aeruginosa in the murine lung and on cystic fibrosis lung airway cells. mBio 3:e00029–12.

27. Bivas-Benita M, Zwier R, Junginger HE, Borchard G. 2005. Non-invasive pulmonary aerosol delivery in mice by the endotracheal route. Eur J Pharm Biopharm 61:214–8.

28. Liu KY, Yang WH, Dong XK, Cong LM, Li N, Li Y, Wen ZB, Yin Z, Lan ZJ, Li WP, Li JS. 2016. Inhalation Study of Mycobacteriophage D29 Aerosol for Mice by Endotracheal Route and Nose-Only Exposure. J Aerosol Med Pulm Drug Deliv 29:393–405.

29. Imamovic L, Serra-Moreno R, Jofre J, Muniesa M. 2010. Quantification of Shiga toxin 2-encoding bacteriophages, by real-time PCR and correlation with phage infectivity. J Appl Microbiol 108:1105–14.

30. Anderson B, Rashid MH, Carter C, Pasternack G, Rajanna C, Revazishvili T, Dean T, Senecal A, Sulakvelidze A. 2011. Enumeration of bacteriophage particles: Comparative analysis of the traditional plaque assay and real-time QPCR- and nanosight-based assays. Bacteriophage 1:86–93.

31. You L, Yin J. 1999. Amplification and spread of viruses in a growing plaque. J Theor Biol 200:365–73.

32. Refardt D. 2012. Real-time quantitative PCR to discriminate and quantify lambdoid bacteriophages of Escherichia coli K-12. Bacteriophage 2:98–104.

33. Duyvejonck H, Merabishvili M, Pirnay JP, De Vos D, Verbeken G, Van Belleghem J, Gryp T, De Leenheer J, Van der Borght K, Van Simaey L, Vermeulen S, Van Mechelen E, Vaneechoutte M. 2019. Development of a qPCR platform for quantification of the five bacteriophages within bacteriophage cocktail 2 (BFC2). Sci Rep 9:13893.

34. Shukla S, Wen AM, Ayat NR, Commandeur U, Gopalkrishnan R, Broome AM, Lozada KW, Keri RA, Steinmetz NF. 2014. Biodistribution and clearance of a filamentous plant virus in healthy and tumor-bearing mice. Nanomedicine (Lond) 9:221–35.

35. Bruckman MA, Randolph LN, VanMeter A, Hern S, Shoffstall AJ, Taurog RE, Steinmetz NF. 2014. Biodistribution, pharmacokinetics, and blood compatibility of native and PEGylated tobacco mosaic virus nano-rods and -spheres in mice. Virology 449:163–73.

36. Zhang YY, Hu KQ. 2015. Rethinking the pathogenesis of hepatitis B virus (HBV) infection. J Med Virol 87:1989–99.

37. Zhang YN, Poon W, Tavares AJ, McGilvray ID, Chan WCW. 2016. Nanoparticle-liver interactions: Cellular uptake and hepatobiliary elimination. J Control Release 240:332–348.

38. Dai J, Chen EQ, Bai L, Gong DY, Zhou QL, Cheng X, Huang FJ, Tang H. 2012. Biological characteristics of the rtA181T/sW172* mutant strain of Hepatitis B virus in animal model. Virol J 9:280.

39. Mostafalou S, Mohammadi H, Ramazani A, Abdollahi M. 2013. Different biokinetics of nanomedicines linking to their toxicity; an overview. Daru 21:14.

40. Nemmar A, Hoet PH, Vanquickenborne B, Dinsdale D, Thomeer M, Hoylaerts MF, Vanbilloen H, Mortelmans L, Nemery B. 2002. Passage of inhaled particles into the blood circulation in humans. Circulation 105:411–4.

41. Inchley CJ. 1969. The actvity of mouse Kupffer cells following intravenous injection of T4 bacteriophage. Clin Exp Immunol 5:173–87.

42. Van Belleghem JD, Clement F, Merabishvili M, Lavigne R, Vaneechoutte M. 2017. Pro- and anti-inflammatory responses of peripheral blood mononuclear cells induced by Staphylococcus aureus and Pseudomonas aeruginosa phages. Sci Rep 7:8004.

43. Van Belleghem JD, Merabishvili M, Vergauwen B, Lavigne R, Vaneechoutte M. 2017. A comparative study of different strategies for removal of endotoxins from bacteriophage preparations. J Microbiol Methods 132:153–159.

44. Speck P, Smithyman A. 2016. Safety and efficacy of phage therapy via the intravenous route. FEMS Microbiol Lett 363.

45. Wienhold SM, Lienau J, Witzenrath M. 2019. Towards Inhaled Phage Therapy in Western Europe. Viruses 11.

46. Raetz CR, Whitfield C. 2002. Lipopolysaccharide endotoxins. Annu Rev Biochem 71:635–700.

47. Rosadini CV, Kagan JC. 2017. Early innate immune responses to bacterial LPS. Curr Opin Immunol 44:14–19.

48. Skirecki T, Cavaillon JM. 2019. Inner sensors of endotoxin - implications for sepsis research and therapy. FEMS Microbiol Rev 43:239–256.

49. Labrie SJ, Samson JE, Moineau S. 2010. Bacteriophage resistance mechanisms. Nat Rev Microbiol 8:317–27.

50. Le S, Yao X, Lu S, Tan Y, Rao X, Li M, Jin X, Wang J, Zhao Y, Wu NC, Lux R, He X, Shi W, Hu F. 2014. Chromosomal DNA deletion confers phage resistance to Pseudomonas aeruginosa. Sci Rep 4:4738.

51. Azam AH, Tanji Y. 2019. Bacteriophage-host arm race: an update on the mechanism of phage resistance in bacteria and revenge of the phage with the perspective for phage therapy. Appl Microbiol Biotechnol 103:2121–2131.

52. Latino L, Midoux C, Vergnaud G, Pourcel C. 2019. Investigation of Pseudomonas aeruginosa strain PcyII-10 variants resisting infection by N4-like phage Ab09 in search for genes involved in phage adsorption. PLoS One 14:e0215456.

53. Lim WS, Ho PL, Li SF-Y, Ow DS-W. 2019. Clinical Implications of Pseudomonas aeruginosa: Antibiotic Resistance, Phage & Antimicrobial Peptide Therapy. Proceedings of the Singapore National Academy of Science 13:65–86.

54. Lin Y, Chang RYK, Britton WJ, Morales S, Kutter E, Chan HK. 2018. Synergy of nebulized phage PEV20 and ciprofloxacin combination against Pseudomonas aeruginosa. Int J Pharm 551:158–165.

55. Chan BK, Sistrom M, Wertz JE, Kortright KE, Narayan D, Turner PE. 2016. Phage selection restores antibiotic sensitivity in MDR Pseudomonas aeruginosa. Sci Rep 6:26717.

